# Modulation of Recombinant Human T-type calcium Channels by Δ^9^-tetrahydrocannabinolic acid *in vitro*

**DOI:** 10.1101/2020.10.11.335422

**Authors:** Somayeh Mirlohi, Chris Bladen, Marina Santiago, Mark Connor

## Abstract

**Introduction:** Low voltage-activated T-type calcium channels (T-type *I*_Ca_), Ca_V_3.1, Ca_V_3.2, and Ca_V_3.3 are opened by small depolarizations from the resting membrane potential in many cells and have been associated with neurological disorders including absence epilepsy and pain. Δ^9^-tetrahydrocannabinol (THC) is the principal psychoactive compound in *Cannabis* and also directly modulates T-type *I*_Ca_, however, there is no information about functional activity of most phytocannabinoids on T-type calcium channels, including Δ^9^-tetrahydrocannabinol acid (THCA), the natural non-psychoactive precursor of THC. The aim of this work was to characterize THCA effects on T-type calcium channels.

**Materials and Methods:** We used HEK293 Flp-In-TREx cells stably expressing Ca_V_3.1, 3.2 or 3.3. Whole-cell patch clamp recordings were made to investigate cannabinoid modulation of *I*_Ca_.

**Results:** THCA and THC inhibited the peak current amplitude Ca_V_3.1 with a *p*EC_50_s of 6.0 ± 0.7 and 5.6 ± 0.4, respectively. 1μM THCA or THC produced a significant negative shift in half activation and inactivation of Ca_V_3.1 and both drugs prolonged Ca_V_3.1 deactivation kinetics. THCA (10 μM) inhibited Ca_V_3.2 by 53% ± 4 and both THCA and THC produced a substantial negative shift in the voltage for half inactivation and modest negative shift in half activation of Ca_V_3.2. THC prolonged the deactivation time of Ca_V_3.2 while THCA did not. THCA inhibited the peak current of Ca_V_3.3 by 43% ± 2 (10μM) but did not notably affect Ca_V_3.3 channel activation or inactivation, however, THC caused significant hyperpolarizing shift in Ca_V_3.3 steady state inactivation.

**Discussion:** THCA modulated T-type *I*_Ca_ currents *in vitro*, with significant modulation of kinetics and voltage dependence at low μM concentrations. This study suggests that THCA may have potential for therapeutic use in pain and epilepsy via T-type channel modulation without the unwanted psychoactive effects associated with THC.

## Introduction

*Cannabis sativa* has been used for thousands of years as a medicinal plant for the relief of pain and seizures^1–3^. There is a growing body of evidence suggesting cannabinoids are beneficial for a range of clinical conditions including pain^4^ inflammation ^5^ epilepsy ^6–8^, sleep disorders^9^, symptoms of multiple sclerosis^10^, and other conditions ^11,12^. Phytocannabinoids, derived from diterpenes in *Cannabis,* have a range of distinct pharmacological actions ^13^. The best characterised phytocannabinoid is Δ^9^-tetrahydrocannabinol (THC), well known for its psychoactive effects ^14^, mediated by its activation of the cannabinoid receptor CB_1_ 15. The next most abundant phytocannabinoid is cannabidiol (CBD), which is non-psychotomimetic and proposed to have potential therapeutic effects in a broad range of neurological disorders ^16–18^ and which has been shown to inhibit signalling via at both CB_1_ and CB_2_ receptors ^16,19,20^ Cannabinoids can also interact with a wide variety of ion channels including Transient Receptor Potential (TRP) channels, ligand gated channels and voltage dependent channels ^21^. THC was identified as a prototypic agonist of TRPA1 and subsequently it and other phytocannabinoids have been reported to activate or inhibit many other TRP channels ^22^. THC and CBD inhibit evoked currents through recombinant 5-HT3 receptors independently of cannabinoid receptors ^23^; and THC caused significant inhibition of native receptor in mammalian neurons ^24^. THC and CBD also potentiate glycine receptor function through an allosteric mechanism^25^.

Voltage gated ion channels also modulated by phytocannabinoids. CBD and cannabigerol (CBG) are able to inhibit voltage-gated Na (Na_V_) channels *in vitro* ^26,27^ which has been suggested to contribute to anti-epileptic effects. A wide range of cannabinoids have been shown to modulate T type *I*_Ca_ channels, including endogenous cannabinoids anandamide and N-arachidonoyl dopamine ^28^, endogenous lipoamino acids such as N-arachidonoyl 5-HT and N-arachidonoyl glycine, as well as the phytocannabinoids THC and CBD ^29–31^. These effects are thought to be mediated by direct interaction of the ligands with channels, as the experiments were done in cells do not express cannabinoid receptors.

Voltage-dependent Ca^2+^ channels are categorized into three families: L-type channels (CaV1), the neuronal N-, P/Q- and R-type channels (CaV2) and the T-type channels (Ca_V_3) ^32^. T-type Ca^2+^ channels (Ca_V_3), can activate upon small depolarizations of the plasma membrane and are present in many excitable cells ^33^ where they are critical for neuronal firing and neurotransmitter release and physiological processes such as slow wave sleep ^34–36^. Cells expressing T-type calcium channels are involved in epilepsy, pain and other diseases and there is substantial evidence supporting the idea that modulating T type calcium channels is a potential therapeutic option in these conditions ^37–39^. T-type calcium are encoded by three Ca_V_3 subunits (Ca_V_3.1, Ca_V_3.2, and Ca_V_3.3). Much smaller membrane depolarizations are required for opening, and at typical neuronal resting membrane potentials a significant number of T-type channels are inactivated. They markedly differ in some of their electrophysiological properties ^40,41^. The most notable of these are that Ca_V_3.1 and Ca_V_3.2 have much faster activation and inactivation kinetics, than Ca_V_3.3 ^42,43^.

Δ^9^-tetrahydrocannabinolic acid (THCA) is the precursor of THC in *Cannabis*. THCA is acutely decarboxylated to form THC by heating^44^. Importantly, THCA has low affinity at CB_1_ receptor ^45^ but interestingly, THCA has been reported to have neuroprotective, anti-inflammatory, and immunomodulatory effects ^44^, raising the possibility of therapeutic activity without unwanted psychotropic effects.

Previous work from our lab have shown that THC and CBD modulate T-type calcium channels ^46^, however, there is no information surrounding the effects of other phytocannabinoids including THCA on these channels. The aim of this work was to characterize THCA modulatory effects on the T-type calcium channels and compare its effects with THC. If THCA could also modulate Ca_V_3 channels, this may provide potential therapeutic activity in pain and other disorders involving the peripheral nervous system without having psychoactive properties.

## Methods

### Transfection and Cell culture

Flp-In T-REx 293 HEK cells (ThermoFisher) were stably transfected with pcDNA5/FRT/TO vector encoding human Ca_V_3.1 (NM 018896.4), Ca_V_3.2 (NM 021098.2), or Ca_V_3.3 (NM 021096.3) (GenScript). The integration of this vector to the Flp-In site was mediated by pOG44, Flp-recombinase expression vector pOG44, which was co-transfected as per manufacturer’s recommendation (ratio 9:1). Transfections were done using Fugene HD transfection agent (Promega) at ratio 1:4 (w/v) total DNA: Fugene HD. Selection of stably expressing cells were performed using 150μg/mL Hygromycin B Gold (InvivoGen) as per kill curve (data not shown). Flp-In T-Rex 293 HEK cells (expressing Ca_V_3.1, Ca_V_3.2, or Ca_V_3.3) do not express CB1 or CB2 receptors^47^. Cells were cultivated in Dulbecco’s modified Eagle’s medium (DMEM) supplemented with 10% FBS, and 1% penicillin-streptomycin. HEK-Ca_V_3.1, Ca_V_3.2, and Ca_V_3.3 were passaged in media with 15μg/ml Blasticidin (InvivoGen) and 100μg/ml Hygromycin. Cells were maintained in 5% CO_2_ at 37°C in a humidified atmosphere. Channel expression was induced by adding 2μg/mL tetracycline.

### Electrophysiology

Currents in Flp-In T-REx 293 HEK cells expressing Ca_V_3.1, Ca_V_3.2, or Ca_V_3.3 channels were recorded in the whole-cell configuration of the patch clamp method at room temperature. Dishes were constantly perfused with external recording solution containing (in mM) (1MgCl_2_, HEPES, 10 Glucose, 114 CsCl, 5 BaCl_2_) (pH to 7.4 with CsOH, osmolarity =330). 2-4 MΩ recording electrodes were filled with internal solutions containing (in mM):126.5 CsMeSO_4_,11 EGTA, 10 HEPES adjusted to pH 7.3 with CsOH. Immediately before use, internal solution was added to a concentrated aliquot of GTP and ATP to yield final concentrations of 0.6 mM and 2mM, respectively. All recordings were measured using an Axopatch 200B amplifier in combination with Clampex 9.2 software (Molecular Devices, Sunnyvale, CA). All data were sampled at 5-10 kHz and filtered at 1 kHz. All currents were leak subtracted using P/N4 protocol.

THC and THCA were prepared daily from concentrated DMSO stocks and diluted in external solution to appropriate concentrations and applied locally to cells via a custom-built gravity driven micro perfusion system. Before running drugs in test of activation and inactivation of Ca_V_3 channels, external control solution was applied about 5 minutes in each experiment to observe in the absence of drugs, vehicle controls itself have no effects on Ca_V_3 channel kinetics. All solutions did not exceed 0.1% DMSO and this concentration of vehicle had no effect on current amplitude or on half activation and half-inactivation potentials (Table1).

**Table 1.** The effects of THCA and THC on the parameters of steady state activation and inactivation of Ca_V_3 channels. Cells expressing recombinant Ca_V_3 channels were voltage-clamped at −100mV then stepped to potential above −75mV(activation) stepped every 5mV. The results of peak currents were fitted to a Boltzmann sigmoidal equation. Changes in the voltage for half activation/inactivation (V_0.5_) of the curve are reported in Table 1. No drug represents time dependent changes under our recording conditions. One-way ANOVA **** indicates *p* value <0.0001, *** indicates *p* value<0.001, **indicates *p* value < 0.01 and *indicates *p* value < 0.02.

| Drug | Cav3 | Change in $V_{0.5}$ | |
| --- | --- | --- | --- |
|  |  | Activation | Inactivation |
| THCA | 3.1 | $-7 \pm 2^{****}$ | $-8 \pm 3^{****}$ |
| THCA | 3.2 | $-4.8 \pm 2^{**}$ | $-6 \pm 2^{***}$ |
| THCA | 3.3 | $3 \pm 1$ | $-4 \pm 1^*$ |
| THC | 3.1 | $-7 \pm 2^{****}$ | $-8 \pm 1^{****}$ |
| THC | 3.2 | $-5 \pm 1^{***}$ | $-9 \pm 2^{****}$ |
| THC | 3.3 | $-1 \pm 0.7$ | $-8 \pm 1^{****}$ |
| No drug | 3.1 | $0.7 \pm 0.2$ | $-1 \pm 0.2$ |
| No drug | 3.2 | $-1 \pm 0.4$ | $-0.6 \pm 0.2$ |
| No drug | 3.3 | $-0.5 \pm 0.2$ | $-1 \pm 0.5$ |

This voltage step was repeated at 12 second intervals (1 sweep) for at least 3 mins to achieve a stable peak *I*_Ca_. Perfusion was then switched to 10μM drugs until maximum inhibition was attained (determined when no more *I*_Ca_ inhibition was observed after 3 successive sweeps). Finally, drug was “washed out” by switching perfusion back to control solution consisting of external buffer with vehicle control.

In order to test whether THCA used contained an appreciable amount of THC, we examined the activity of THCA in a fluorescent assay of CB1-dependent activation of inwardly rectifying K channels (described in detail in ^48^). In these experiments, THC (1μM) produced a change in fluorescence of 12.8 ± 1.2 %. In parallel experiments, THCA (1μM) did not significantly alter the fluorescence (1.0 ± 0.6%). *p*EC_50_ for THC in this assay is about 300nM ^48^ and 100nM THC produces a robust change in fluorescence ^49^, the lack of effect of THCA at 1μM suggests that there was no significant contamination of THCA with THC.

### Drugs and reagents

The THC and THCA used in this study were a kind gift from University of Sydney’s Lambert Institute for Cannabinoid Therapeutics. Drugs (30 mM) were aliquoted and stored as concentrated stocks in DMSO and stored at −30 C. Daily dilutions were made fresh before each use in external recording solution to give a final vehicle concentration of 0.1%.

### Statistics

Data are reported as the mean and standard error of at least 6 independent experiments. Concentration response curves, steady state inactivation and activation were generated by fitting data to a Boltzmann sigmoidal equation in Graph Pad Prism 8. Statistical significance for comparing the V_0.5_ values of activation and inactivation were determined using one-way ANOVA comparing values of V_0.5_ calculated for individual experiments. In order to compare the changes in the time to peak and decay time of deactivation, unpaired t-test was used. All values are reported as mean ± standard errors and were fitted with a modified Boltzmann equation: I = [Gmax*(Vm-Erev)]/[1+exp((V_0.5_ act-Vm)/ka)], where V_m_ is the test potential, V_0.5_ act is the half-activation potential, Erev is the reversal potential and Gmax is the maximum slope conductance. Steady-state inactivation curves were fitted using Boltzmann equation: I =1/ (1 + exp ((Vm - Vh)/k)), where Vh is the half-inactivation potential and k is the slope factor.

## Results

Superfusion of THCA and THC on Ca_V_3 inhibited the peak of the *I*_Ca_ evoked by a step from −100mV to −30 mV (Fig 1). At a concentration of 10 μM, THC or THCA blocked the current amplitude of Ca_V_3.1 almost completely, and inhibited Ca_V_3.2 by 56 ± 2% and 53 ± 4 % respectively (n=6). 10μM THC did not affect Ca_V_3.3 *I*_Ca_ while 10μM THCA inhibited Ca_V_3.3 by 43% ± 2 (Fig 1A). Ca_V_3.1 was inhibited by THC and THCA with *p*EC_50_ 6 ± 0.7 and 5.6 ± 0.4 respectively (Fig1B). The effects of THCA and THC on Ca_V_3.1, 3.2 and 3.3 currents are illustrated in Fig 2 (THCA) and Fig 3 (THC), the drug effects did not readily reverse on washout.

**Figure 1.**
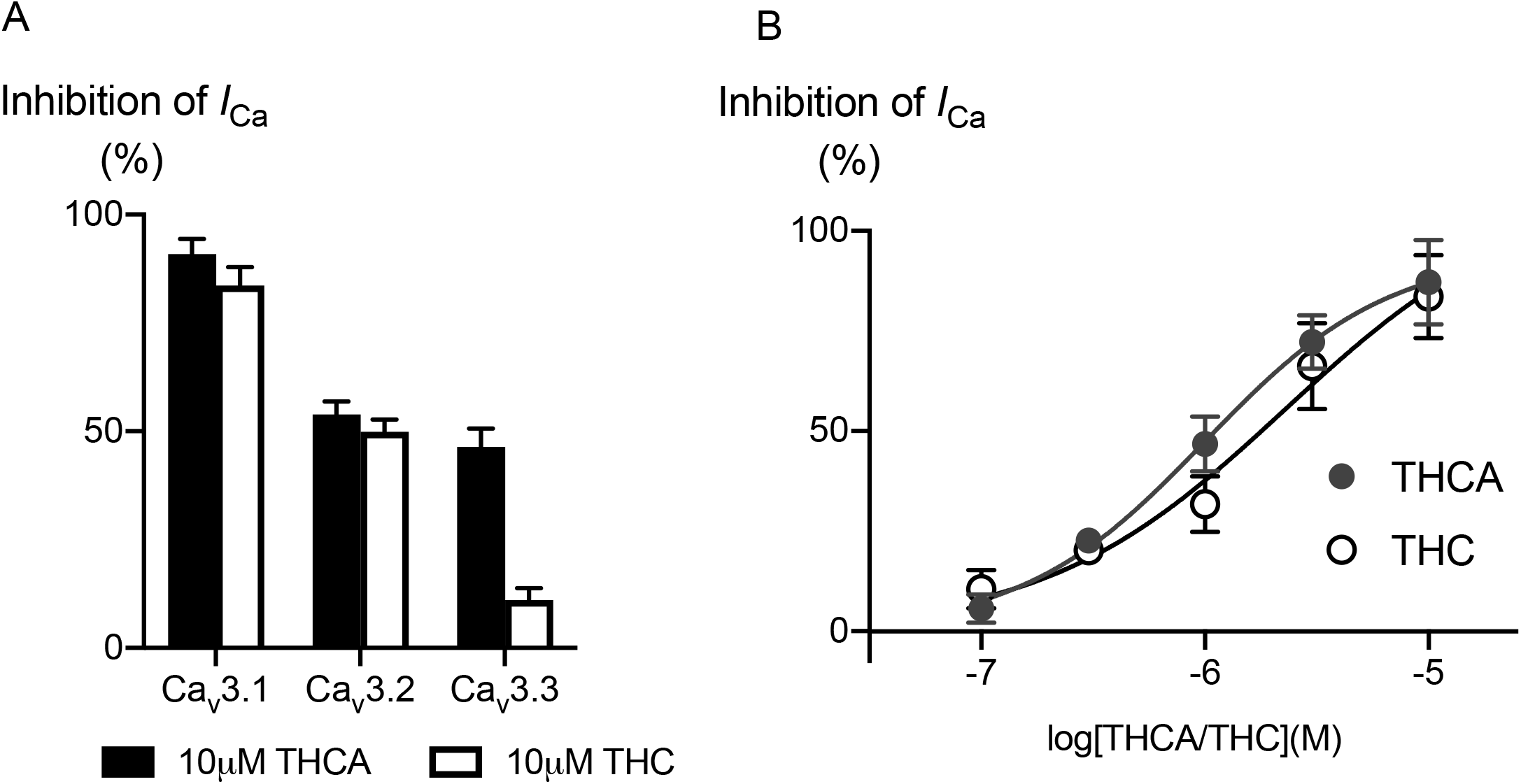
Effects of 10μM THCA and THC on T-type calcium channel current and concentration response curve for THCA and THC effects on Ca_V_3. **(A)** Peak *I*_Ca_ was elicited by a step from −100 mV to −30 mV; Ca_V_3.1 current was almost completely inhibited by10 μM application of THC and THCA. 10 μM THC and THCA blocked Ca_V_3.2 calcium current about 52% ±3. Ca_V_3.3 current was not affected by10 μM THC but THCA decreased Ca_V_3.3 calcium current by 43% ± 4. **(B)** Concentration response curves was created to determine the potency of these compounds at Ca_V_3.1. Each point represents the mean ± SEM of 6 cells.

**Figure 2.**
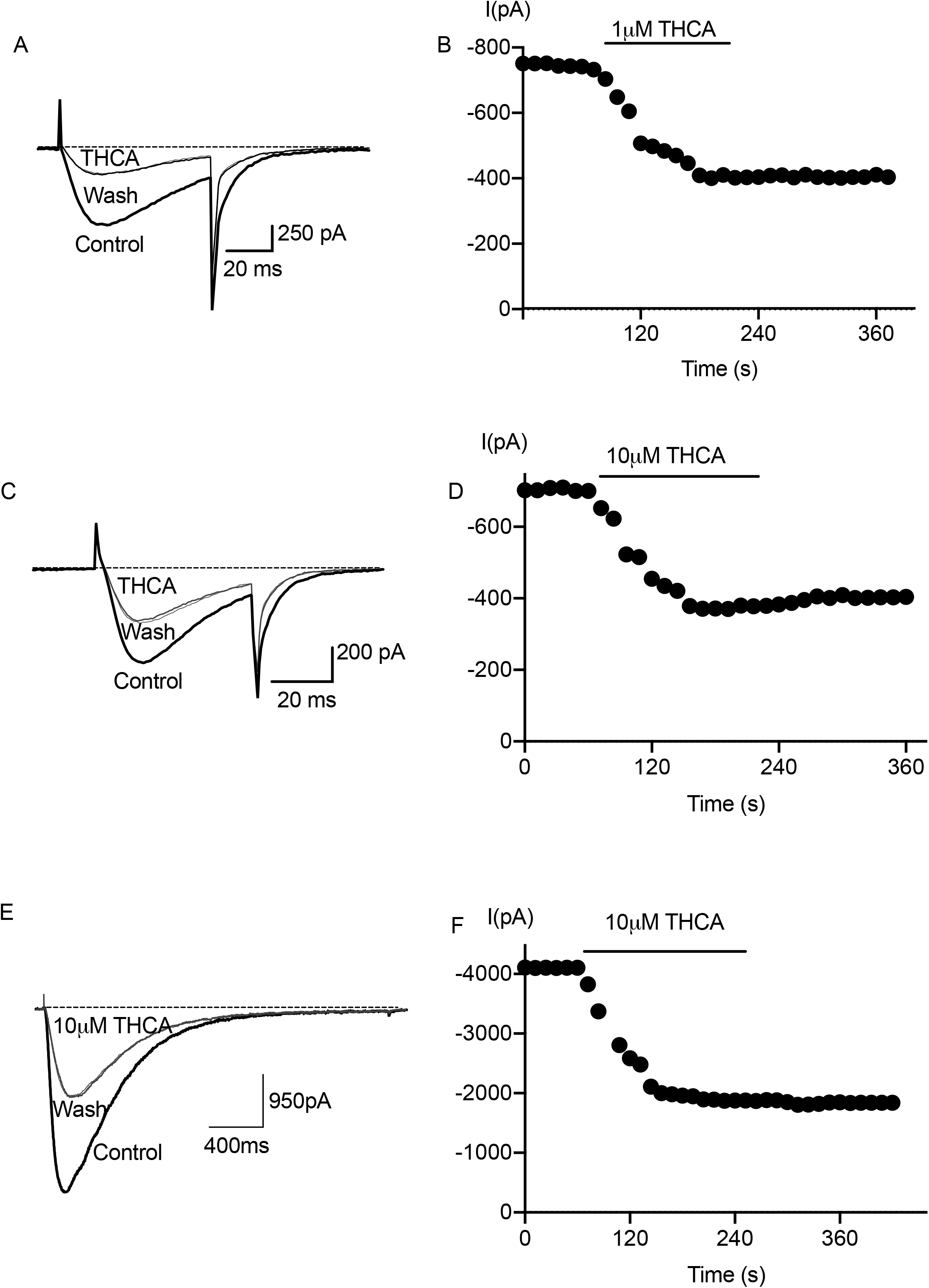
THCA effects on Ca_V_3 current amplitude. Each trace represents the current elicited by a voltage step from −100 mV to −30 mV. **(A)** 1μM THCA inhibited calcium current of Ca_V_3.1. **(B)** Time course of inhibition and degree of reversibility THCA inhibition of Ca_V_3.1 is illustrated. **(C)** THCA 10μM inhibited calcium current of Ca_V_3.2. **(D)** Time course of inhibition and degree of reversibility THCA inhibition of Ca_V_3.2 is illustrated. **(E)** THCA at 10 μM inhibited current amplitude of Ca_V_3.3. **(F)** The inhibition of Ca_V_3.3 by 10 μM THCA was not washed out shown in time course inhibition of Ca_V_3.3.

**Figure 3.**
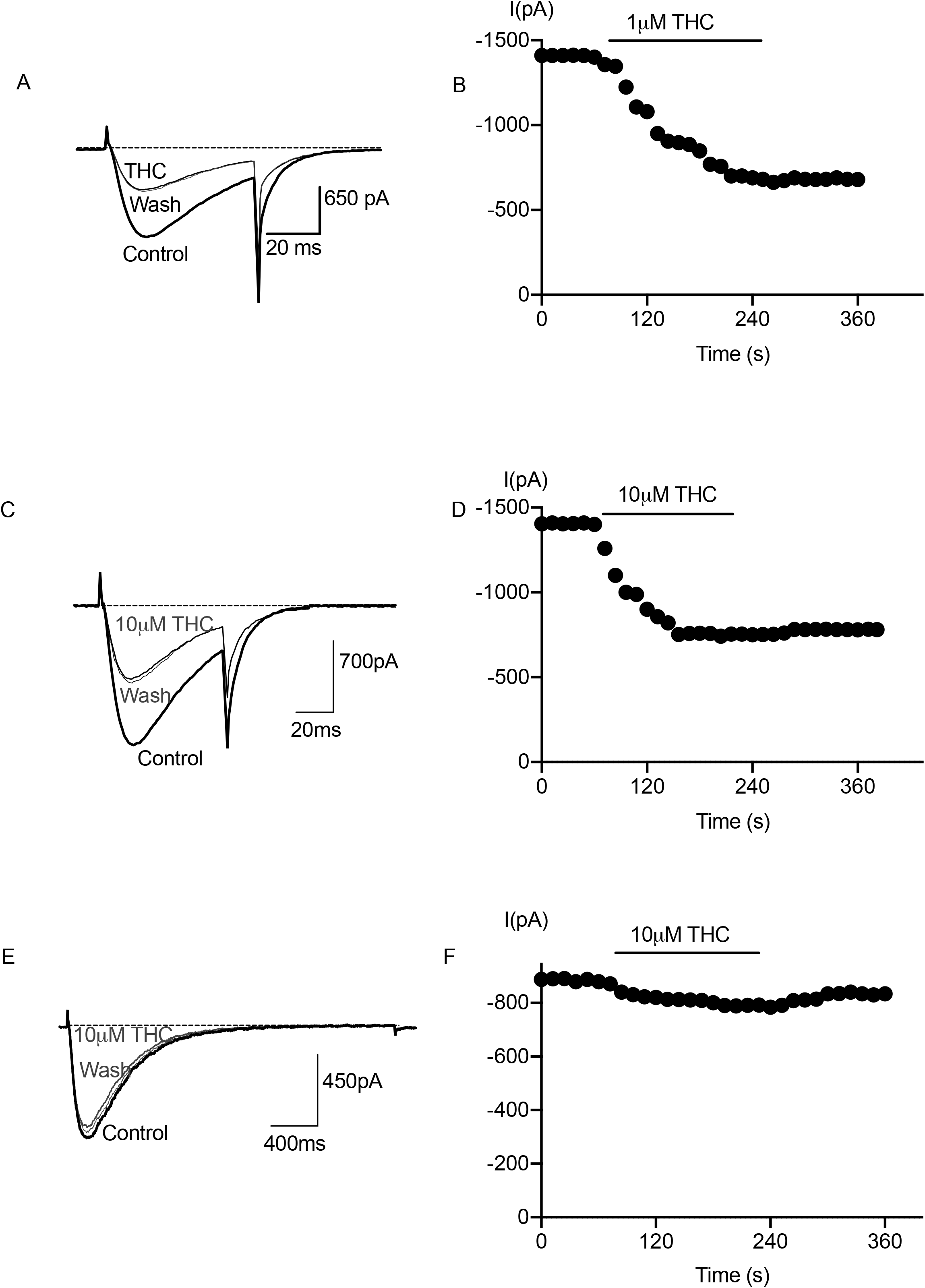
THC effects on Ca_V_3 current amplitude. Recording of Ca_V_3 channel was made as outlined under experimental procedures. Each trace represents the current elicited by a voltage step from −100mV to −30mV. **(A)** 1 μM THC inhibited Ca_V_3.1 calcium current. **(B)** inhibitory effects of THC on Ca_V_3.1 was not washed out by using external solution. **(C)** THC inhibited Ca_V_3.2 calcium current at 10 μM. **(D)** A reversal of THC (10 μM) inhibition of Ca_V_3.2 was not seen by washing. **(E)** THC at 10 μM had little effect on calcium current of Ca_V_3.3. **(F)** Inhibition by THC at 10 μM was not reversible.

### THC and THCA effects on activation and inactivation kinetics

We examined the voltage-dependence of activation Ca_V_3 channels by repetitively stepping cells from −75mV to 50mV from a holding potential of −100mV. After a control I/V relationship was generated, it was repeated after 5 min perfusion of THCA (Fig 4A). The voltage-dependence of activation for Ca_V_3.1 was affected by THCA, notably it increased current amplitudes for depolarisations between −75mV to −45mV and inhibited current amplitude for depolarisations between −35 and 50mV (Fig 4B). THCA produced a significant hyperpolarizing shift in the half activation potential of Ca_V_3.1; these shifts were not seen with time-matched vehicle controls (Table 1). Steady-state inactivation, where cells were voltage clamped at potentials between (−110 mV and −20 mV) for 2s before current were evoked by stepping them to test potentials of −30mV, showed that THCA also caused large shifts in steady-state inactivation of Ca_V_3.1 (Fig 4C). Activation and inactivation changes for cells exposed to vehicle alone for 5 min were less than −1mV (Table1). Using the same protocols, it was found that THCA also shifted Ca_V_3.2 half activation to negative potentials and caused a larger shift in half inactivation of Ca_V_3.2 (Fig 4D). THCA caused small positive shift and significant negative shift in half activation and inactivation of Ca_V_3.3 (Fig 4E).

**Figure 4.**
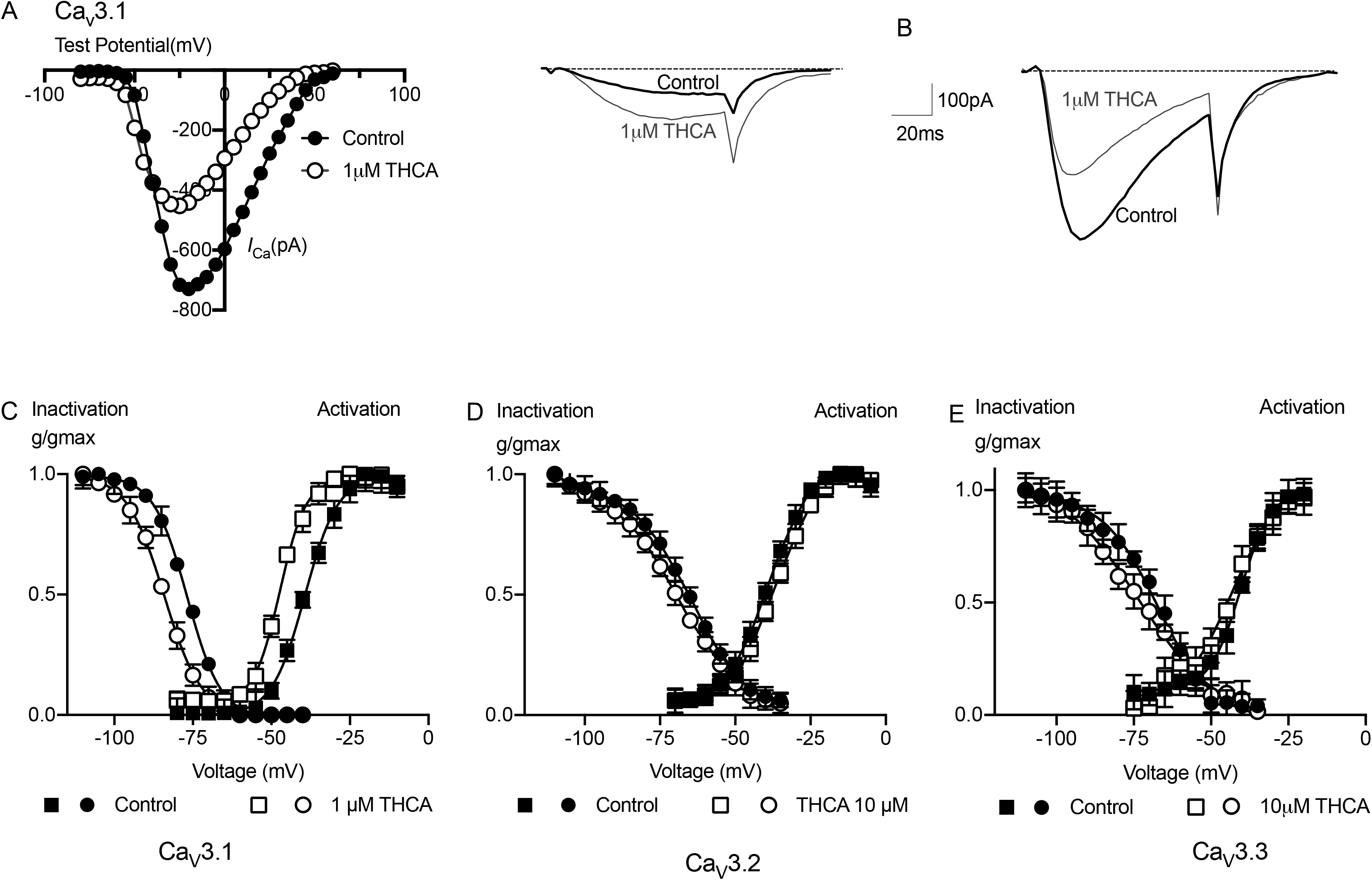
THCA effects on the activation and inactivation of Ca_V_3 channels. **(A)** Current-Voltage (I-V) relationship showing the activation of Ca_V_3.1 from a holding membrane potential of −100mV in the absence and presence of 1μM THCA. The peak current amplitude is plotted, **(B)** example traces of this experiment illustrating the effects of 1μM THCA at testing membrane potential of −51mVand −22mV: current is enhanced at lower test potentials then inhibited at more depolarized potentials. **(C)** 1 μM THCA affected half activation and inactivation of Ca_V_3.1 expressed in HEK293 to negative potentials. **(D)** Steady state activation and inactivation of Ca_V_3.2 expressed in HEK293 in the presence and absence of THCA showed a significant shift in inactivation of Ca_V_3.2 however 10 μM THCA created slight shift in activation of Ca_V_3.2. **(E)** THCA caused a small positive shift in activation kinetics of Ca_V_3.3 and a small negative shift in inactivation of Ca_V_3.3. Each data points represent the mean ± SEM of 6 cells.

1μM THC also affected steady state inactivation and activation of Ca_V_3.1. THC shifted half activation and inactivation of Ca_V_3.1 to more negative voltages (Fig 5C). THC shifted half activation of Ca_V_3.2 to negative potentials and caused significant negative shift in inactivation of Ca_V_3.2 (Fig 5D). THC at 10μM had no effect on the half activation of Ca_V_3.3 however THC negatively shifted the half inactivation of Ca_V_3.3 significantly (Fig 5E).

**Figure 5.**
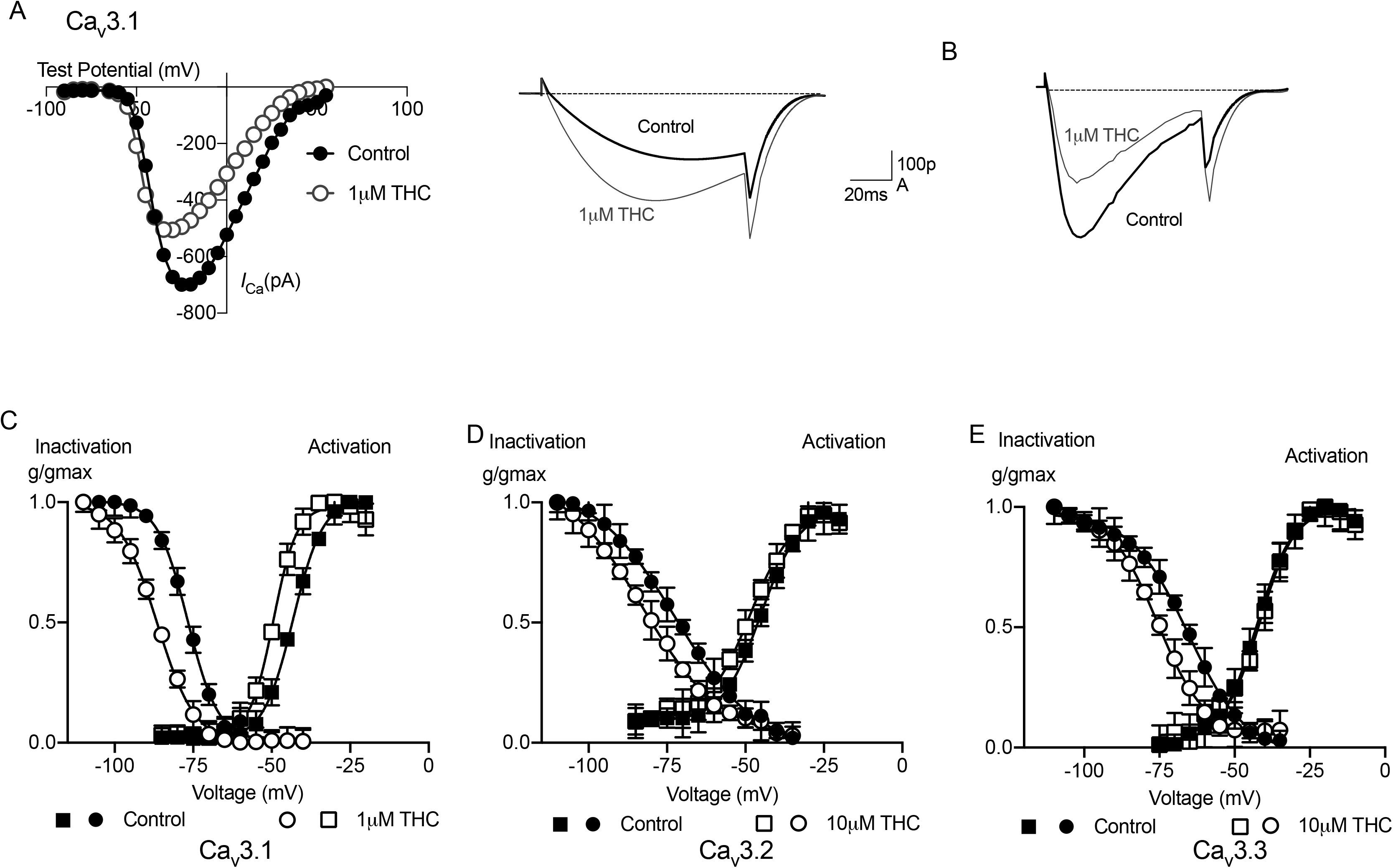
Effects of THC on the voltage-dependence of Ca_V_3 activation and Inactivation. Current-Voltage (I-V) relationship showing the activation of Ca_V_3.1 from a holding membrane potential of −100mV in the absence and presence of 1 μM THC.**(B)** The peak current amplitude is plotted at testing membrane potential of −51mV and −22mV. Example traces of this experiment illustrating the effects of 1μM THC: current is enhanced at lower test potentials then inhibited at more depolarized potentials.**(C)** THC effect on Ca_V_3 channels kinetics when HEK293 cells were voltage clamped at −100mV, depolarized to 50mV from −75mV showed that 1 μM THC shifted activation and inactivation of Ca_V_3.1 to negative potentials significantly.**(D)** 10 μM THC effects on activation and inactivation kinetics of Ca_V_3.2 indicated steady state inactivation was shifted to negative potentials significantly however THC caused −5mV shift in activation kinetics of Ca_V_3.2.**(E)** 10 μM THC effects on activation and inactivation kinetics of Ca_V_3.3; THC had no effects on steady state activation however THC caused significant shift in inactivation kinetics of Ca_V_3.3. Each data point represent the mean ± SEM of six cells.

### Effects of THC and THCA on time to peak and kinetics of current deactivation of Ca_V_3 channels

THC and THCA caused no significant changes on time to peak on any of the T-type channels at any voltage (Fig 6A-F). The effects of THC and THCA on deactivation of currents elicited during the standard I/V protocol, were measured by fitting a monophasic exponential to the inward “tail” currents that resulted immediately following the voltage step. 1μM THCA slowed deactivation of Ca_V_3.1 (Fig. 7A, C), however, the deactivation of both Ca_V_3.2 (Fig 7E) and Ca_V_3.3 (not shown) were unaffected by THCA at 10μM. THC slowed deactivation of Ca_V_3.1 (1 μM, Figure 7B, D) and Ca_V_3.2 (10 μM, Figure 7F) but THC did not change deactivation of Ca_V_3.3 (not shown).

**Figure 6.**
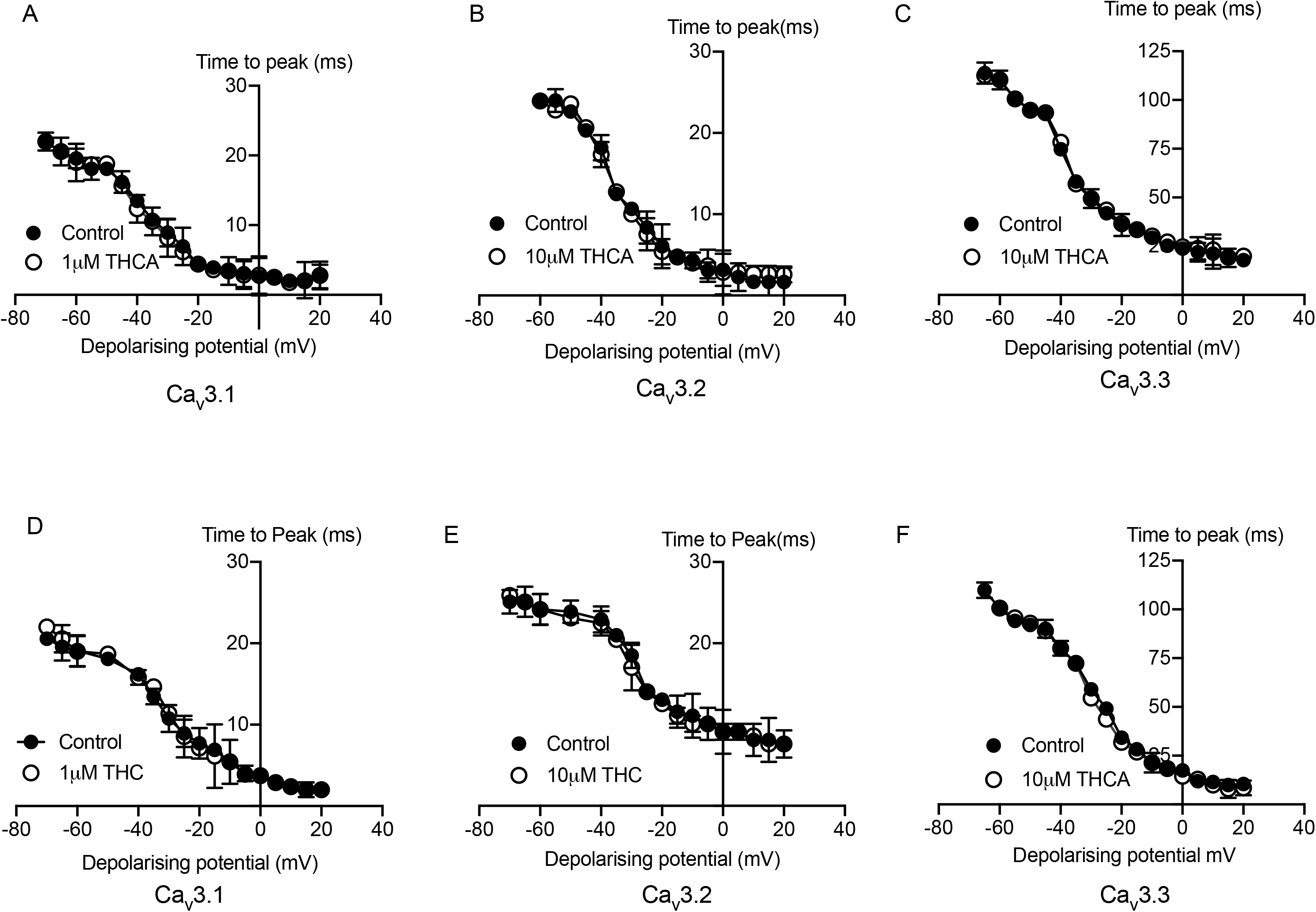
THCA and THC effects on time to peak of Ca_V_3 channels. The plots illustrate the time to peak of current Ca_V_3 before and after 5min superfusion of THC and THCA. THCA had no significant effects on time to peak of **(A)** Ca_V_3.1, **(B)** Ca_V_3.2 and **(C)** Ca_V_3.3. No shift was seen to those in parallel THC experiments where solvent alone was super fused for **(D)** Ca_V_3.1, **(E)** Ca_V_3.2 and **(F)** Ca_V_3.3. Each point represents the mean ± SEM of at least six cells (Unpaired t-test P>0.05).

**Figure 7.**
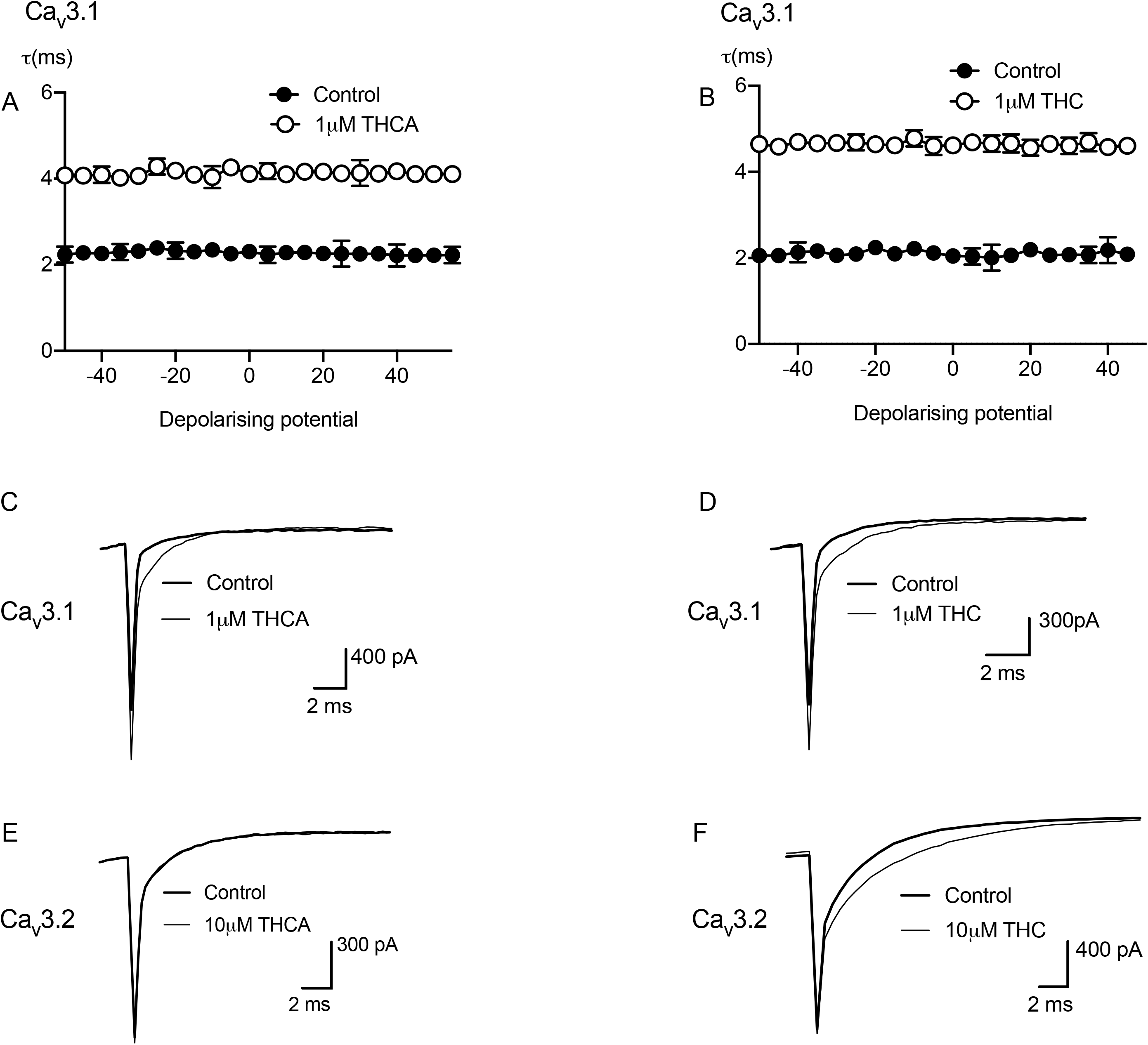
THCA and THC effects on Ca_V_3 time constant of deactivation. Cells expressing Ca_V_3 channels were stepped repetitively from a holoing potential of −100 mV to test potentials between −75 and 50 mV. **(A)** THCA produced a significant change in time constant deactivation of Ca_V_3.1(ANOVA, P < 0.0001) across a range of potential membrane. **(B)** THC produced significant changes in time constant deactivation of Ca_V_3.1 across a range of membrane potential (ANOVA, P < 0.0001). **(C)** 1μM THCA prolonged deactivation of Ca_V_3.1 showing in example trace of tail current from I-V current relationships. **(D)** Example traces of tail current for Ca_V_3.1 showed that 1μM THC slowed deactivation of Ca_V_3.1. **(E)** Representative traces illustrated that THCA at 10μM did not affect Ca_V_3.2 and **(F)** 10μM of THC slowed deactivation of Ca_V_3.2.

## Discussion

The major finding of this study is that THCA inhibited T-type calcium channels with most potent effects on Ca_V_3.1. THC also most potently affected Ca_V_3.1, and Ca_V_3.2 was moderately inhibited by both drugs at 10μM with less inhibition of Ca_V_3.3. THCA shifted the half activation and inactivation voltages of Ca_V_3.1 and Ca_V_3.2 to more negative potentials, THC behaved in a similar fashion. THCA and THC also slowed the time constant deactivation of Ca_V_3.1 however at 10μM only THC slowed the deactivation of Ca_V_3.2. Both THCA and THC produced modest shifts in Ca_V_3.3 inactivation without any effects on the deactivation kinetics. The presence of the carboxylic acid moiety in THCA does not result in substantial differences in modulation of T type calcium channel compared with THC.

THC has higher affinity to cannabinoid receptors CB_1_ and CB_2_ ^15^ and causes a distinctive intoxication via activation of the CB_1_ ^50^ receptors, however, studies of affinity of THCA for the CB_1_ receptor have produced different results, but studies where THCA was tested for THC produced by THCA degradation, there was little activity attributable to THCA^51,52^. Verhoeckx *et al* examined THC and THCA affinity using radioligand binding assay and determined that THC had greater affinity compared to THCA at CB_1_^44^. However, Ahmed *et al* reported no affinity of THCA on CB_1_^53^ while Husni *et al.,* found some activity on CB_1_ ^54^, while the one study that reported THC and THCA had similar affinity for CB_1_, did not examine the potential contamination of THCA with THC^51^. We tested the activity of our THCA in a membrane potential assay in AtT20 cells expressing CB_1_ receptors. THC (1 μM) produced a significant hyperpolarization of the cells, as reported many times previously, while THCA did not produce changes in fluorescence, suggesting that in our experiments, THC contamination of the THCA was insignificant.

In current study, THCA like THC shifted steady sate inactivation of the Ca_V_3.1 and Ca_V_3.2 channels to more negative potentials, reducing the number of channels that can open when the cell is depolarised, preventing their transition to an inactivated state. THCA had the same effect as THC on Ca_V_3.1 steady state activation, causing a hyperpolarising shift so that when the cells are depolarised, more channels are available for activation. THCA effects on Ca_V_3.2 kinetics were less pronounced than THC, causing a more negative shift in both activation and inactivation of Ca_V_3.2. The effects of THCA and THC on in half activation of Ca_V_3.3 was not significant. Conversely, THCA and THC caused a significant shift in steady state inactivation of Ca_V_3.3. Interestingly, THCA and THC potentiated Ca_V_3.1 current evoked by modest depolarization and then inhibited current amplitudes following stronger depolarisation. These data suggest that THCA and THC may increase the initial depolarizing drive produced by Ca_V_3.1 in some circumstances, despite the overall inhibitory effects on the channels.

The results with THC are in good agreement with previous studies from our lab. In general, THC showed modestly higher potency to inhibit Ca_V_3.2 and Ca_V_3.3 in the study of Ross *et al*, this can be attributed to the subtle different recording conditions where potency was determined in cells voltage clamped at slightly more depolarized potentials (−100mV vs −86mV)^28^.

Both THC and THCA have been reported to activate TRPA1 and TRPV2 channels and showed the similar antagonist activity on TRPV1 and TRPM8^21,22,55^. Together with the results of our study, these data show that THCA and THC generally behave in a similar manner for ion channel modulation, but they have very different activity on cannabinoid GPCR. The very limited permeability of THCA to cross the blood brain barrier suggests a potential role as a drug for treatment of pain and inflammation in the periphery, and THCA has been shown to reduce inflammation in the gut ^57^. While the mechanism(s) underlying this are still unknown, inhibition of T-Type *I*_Ca_ is a possible contributor. ^58,59^

## ABBREVIATIONS

*I*_Ca_: Voltage gated calcium channel current
THC: Δ^9^-tetrahydrocannabinol
THCA: Δ^9^-tetrahydrocannabinolic acid
CBD: Cannabidiol
TRP: Transient Receptor Potential

## Acknowledgment

We would like to thank Lambert initiative for gift of THC and THCA. We would also like to thank Shivani Sachdev for performing some of the experiments with THCA and THC on CB1 receptor signalling.

